# *In-situ* serial crystallography facilitates 96-well plate structural analysis at low symmetry

**DOI:** 10.1101/2024.04.28.591338

**Authors:** Nicolas Foos, Jean-Baptiste Florial, Mathias Eymery, Jeremy Sinoir, Franck Felisaz, Marcus Oscarsson, Antonia Beteva, Matthew W. Bowler, Didier Nurizzo, Gergely Papp, Montserrat Soler-Lopez, Max Nanao, Shibom Basu, Andrew A. McCarthy

## Abstract

The advent of serial crystallography has rejuvenated and popularised room temperature X-ray crystal structure determination. Structures determined at physiological temperature reveal protein flexibility and dynamics. In addition, challenging samples (e.g., large complexes, membrane proteins, and viruses) forming fragile crystals, are often difficult to harvest for cryo-crystallography. Moreover, a typical serial crystallography experiment requires a large number of microcrystals, mainly achievable through batch crystallisation. Many medically relevant samples are expressed in mammalian cell-lines, producing a meagre quantity of protein that is incompatible for batch crystallisation. This can limit the scope of serial crystallography approaches. Direct *in-situ* data collection from a 96-well crystallisation plate enables not only the identification of the best diffracting crystallisation condition, but also the possibility for structure determination at ambient conditions. Here, we describe an *in situ* serial crystallography (iSX) approach, facilitating direct measurement from crystallisation plates, mounted on a rapidly exchangeable universal plate holder deployed at a microfocus beamline, ID23-2, at the European Synchrotron Radiation Facility (ESRF). We applied our iSX approach on a challenging project, Autotaxin, a therapeutic target expressed in a stable human cell-line, to determine a structure in the lowest symmetry *P*1 space group at 3.0 Å resolution. Our *in situ* data collection strategy provided a complete dataset for structure determination, while screening various crystallisation conditions. Our data analysis reveals that the iSX approach is highly efficient at a microfocus beamline, improving throughput and demonstrating how crystallisation plates can be routinely used as an alternative method of presenting samples for serial crystallography experiments at synchrotrons.

**Synopsis:** The determination of a challenging structure in the *P*1 space group, the lowest symmetry possible, shows how our *in-situ* serial crystallography approach expands the application of crystallisation plates as a robust sample delivery method.

## 1. Introduction

Conventional macromolecular crystallography at cryogenic temperature (100K) (cryo-MX) has so far been the silver bullet in determining structures of biomolecules at near-atomic resolution. This has been complemented with the advent of “resolution revolution” in cryo-electron microscopy (cryo-EM) (Kuhlbrandt, 2014; Amunts *et al*., 2014). However, structure determination at cryogenic conditions does not reveal biomolecular flexibility in native conformations (Fraser *et al*., 2011; Fischer, 2021). Moreover, it does not permit the study of time-dependent structural changes coupled with enzymatic catalysis or protein dynamics (Keedy *et al*., 2015; Weinert *et al*., 2017; Doukov *et al*., 2020; Fischer, 2021). Room temperature crystallography (RT-MX) was initially proposed as a potential solution to this problem. In early days, RT-MX was prone to high radiation damage and limited resolution due to the very low deposited dose allowed. Recent advances in serial crystallography at X-ray free electron lasers (XFELs) and synchrotrons have rejuvenated RT crystallography (Chapman *et al*., 2011; Diederichs & Wang, 2017; Pearson & Mehrabi, 2020). Recent studies revealed that RT-MX performed with *in-situ* data collection strategies has the potential to identify physiological conformations (Fraser *et al*., 2011), allosteric networks (Keedy *et al*., 2018), ligand binding modes (Le Maire *et al*., 2011; J. Gildea *et al*., 2022), and fragment screening (Huang *et al*., 2022). Moreover, RT-MX avoids the hassle of freezing crystals - which can be challenging for membrane proteins, viruses, or large complexes while still yielding atomic resolution (Gavira *et al*., 2020). Therefore, RT-MX has tremendous potential to improve structure-based drug discovery by facilitating a shortest path from protein to physiologically relevant structures. To this end, synchrotron facilities have already enabled RT data collection either through 96 well-plates (Le Maire *et al*., 2011; Axford *et al*., 2012; Bingel-Erlenmeyer *et al*., 2011; Doukov *et al*., 2020), loops with sleeves (Russi *et al*., 2017; Bowler *et al*., 2015), *in-meso in-situ* COC chips for membrane proteins (Huang *et al*., 2015, 2018), microfluidic chip (De Wijn *et al*., 2021), serial synchrotron crystallography (Weinert *et al*., 2017; Diederichs & Wang, 2017) or even dedicated beamlines for *in-situ* experiments e.g., VMXi at the Diamond Light Source (DLS) (Mikolajek *et al*., 2023). RT-MX with *in-situ* data collection is typically executed by measuring small wedges (10° – 60°) from single-crystals in 96 well-plates (Mikolajek *et al*., 2023; Russi *et al*., 2017). This requires visually inspecting each of 96 drops, optical focusing with an on-axis camera, and finally collecting diffraction data from selected positions (dependent on the drop position in the plate). All-together, one 96 well-plate can easily consume 6 – 8 hrs (i.e., a single-shift of allocated beamtime). Thus, the traditional *in-situ* data collection lacks high-throughput and efficiency, which in turn discourages users from routine usage. This limits the application of *in-situ* data collection to only diffraction-based screening of crystallisation conditions of challenging projects. Here, we describe the first implementation of a plate gripper at a microfocus MX beamline – ID23-2 (Nanao *et al*., 2022) at the European Synchrotron Radiation Facility (ESRF), enabling an easy setup, fast switching to and from a standard cryo-setup, and rapid data collection by importing visual scores of drops executed in advance. We transformed *in-situ* data collection into a serial crystallography approach by raster scanning each drop. Our so-called *in-situ* serial crystallography (iSX) has been applied for the first time to determine a challenging structure in the *P*1 space group directly from a CrystalDirect^TM^ plate (Cipriani *et al*., 2012; Márquez & Cipriani, 2014) in order to establish the power of this approach. Thereby, *in-situ* serial crystallography (iSX) directly on 96-well plates will surmount the current *status-quo* of plate application by providing a complete dataset as a by-product.

In this work, autotaxin from rat (r-ATX), which is a phosphodiesterase producing the lipid signalling molecule lysophosphatidic acid (LPA), has been used as a challenging case study. ATX is the main producer of LPA, forming the ATX-LPA signalling axis with LPA acting as a multifunctional lipid mediator by engagement with six dedicated G-protein coupled receptors (LPA_1-6_) (Eymery *et al*., 2023). ATX regulates many pathophysiological processes e.g., vascular development, neuropathy pain, fibrosis, rheumatoid arthritis, sclerosis, and cancer (Moolenaar & Perrakis, 2011). The X-ray structure of r-ATX (MW 100 kDa) has already been determined through conventional cryo-MX (Hausmann *et al*., 2011; Eymery *et al*., 2023, 2024), giving a good control over crystallisation, which was minimally optimised to obtain sitting drops containing micro-crystals. r-ATX was expressed in a HEK293T cell-line, producing ∼4 mg/mL purified protein (Eymery *et al*., 2023), which is inadequate for batch crystallisation. Thus, r-ATX serves as a relevant real-life case, justifying the viability of a crystallisation plate as the most natural sample delivery approach for serial crystallography experiments. Here, we show how the application of iSX on micro-crystals facilitated the determination of the first room temperature structure of r-ATX at 3.0 Å in the *P*1 space group with an effective data collection of < 1.5 hrs.

## 2. Experimental setup at ID23-2 beamline

ID23-2, a true microfocus MX beamline, was completely rebuilt (Nanao *et al*., 2022) to exploit the benefits of the extremely brilliant source (EBS) upgrade at the ESRF (Raimondi *et al*., 2023). The beamline provides a fixed energy (14.2 keV) with variable micro-focusing from 2 – 10 µm^2^ beam size at a photon flux of 1.3 x 10^13^ photons/sec at 200 mA ring-current. ID23-2 is equipped with a high-precision MD3-Up diffractometer (Arinax, Moirans, France) and EIGER2 9M detector (Casanas *et al*., 2016). The experimental setup for *in-situ* data collection uses high-precision features of the standard MD3-Up goniometer, complemented by an additional horizontal axis for aligning the crystallisation plate on the omega axis of rotation. An attractive feature of the MD diffractometers is exchangeable goniometer-heads. For cryo-MX, we typically use a mini-kappa 3 (MK3; (Brockhauser *et al*., 2013)) goniometer-head (**Figure 1**), which can easily be switched for a dedicated plate-holder, so-called Plate Manipulator (PM) for mounting SBS compatible 96-well plates in < 15 min (**Figure 1**). Both goniometer-heads – MK3 and PM utilise the same underlying translational stages to centre the crystal with the on-axis camera. In brief, given that the rest of MD3-Up diffractometer retains the same functionality, the switching of portable goniometer-heads allows for a rapid change from a cryo-MX to *in situ* setup.

**Figure 1.**
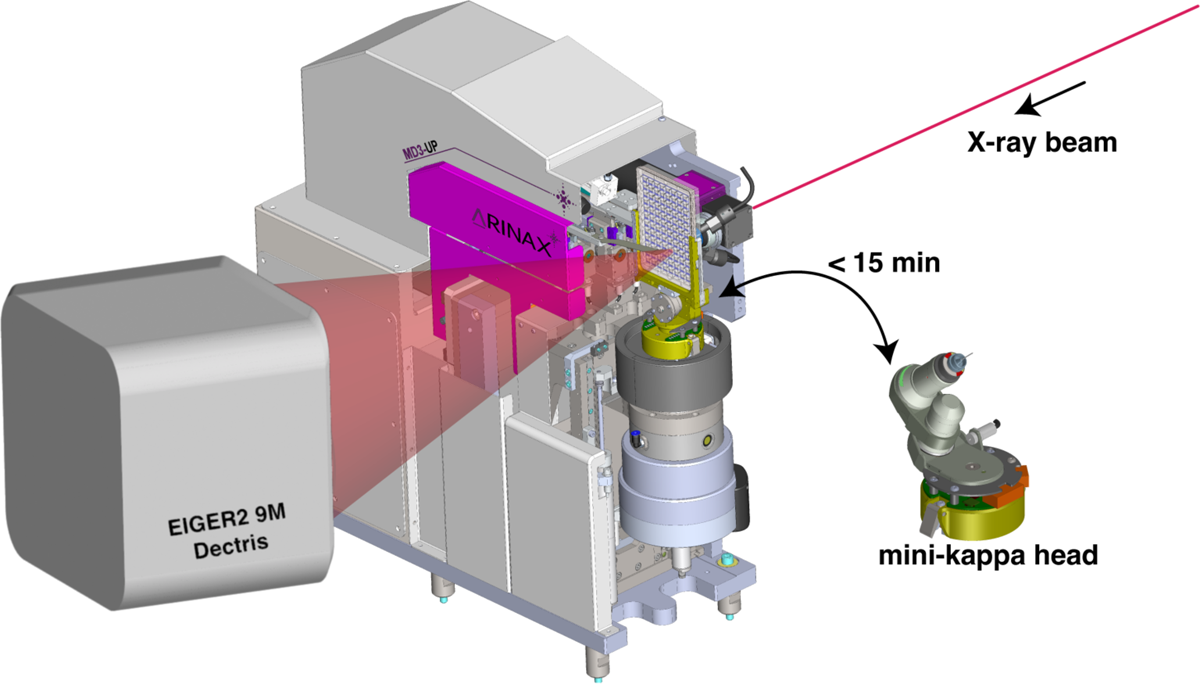
A schematic for the *in-situ* experimental setup on ID23-2 with the Plate Manipulator installed on a MD3-Up and a CrystalDirect plate loaded. It also illustrates the fast switching from the conventional mini-kappa goniometer-head to *in-situ* setup (Diffractometer drawing kindly provided by Arinax, Moirans, France).

## 3. Integration of the plate-holder at ID23-2 beamline

### 3.1 Plate manipulator for Micro-diffractometer

The Plate Manipulator (PM) is a hardware device with a fork shaped design (**Figure 2a**) that is compatible with the following SBS plates: In Situ-1, Mitegen; Crystal-Direct, Mitegen; and CrystalQuick-X, Greiner Bio-One (**Figure 2b**). The PM horizontal axis for navigating on the plate relies on two different types of motions. The long-range motion is executed by a cable transmission driven by a stepper motor, while the short-range motion is performed by the centring-table stage of the micro-diffractometer. The centring-table is highly precise with < ± 2 µm within a 10 mm range and the effective resolution for crystal alignment is 300 nm. The vertical motion is performed by the MD3-Up high precision vertical translation axis. The rotation capability has not been used in our protocol but is available through the omega rotation axis. The accessible omega range is up to 70° depending on the well location in the plate and beamstop distance. Data collection can be performed over the whole area of the SBS plate regardless of the well position. It also allows the use of different data-collection procedures, using mesh, single or multi-point as well as pure raster scanning with a large range of oscillation. Similar to the *in-situ* implementation on ID30B (McCarthy *et al*., 2018), in order to exchange from the MK3 to the PM setup, the cryo-stream and beam-cleaning capillary are removed and disabled, while the beam-defining aperture is replaced with a 20 mm long and 100 µm diameter canon aperture (a combined aperture and beam-cleaning capillary). Both goniometer-heads at ID23-2 benefit from the latest version of the ‘quick lock’ mechanism to facilitate a rapid and reproducible positioning on the omega axis. A number of interlocks specific to the PM head are auto-configured to prevent potential collisions with diffractometer organs (e.g., backlight and beamstop). It takes ∼15 min to exchange goniometer heads and configure the MD3-Up for *in situ* data collection. The crystallisation plates are manually mounted. However, remote experiments can be carried out upon request. Executing a MeshScan over a 4 x 4 mm^2^ area requires ∼10 min.

**Figure 2.**
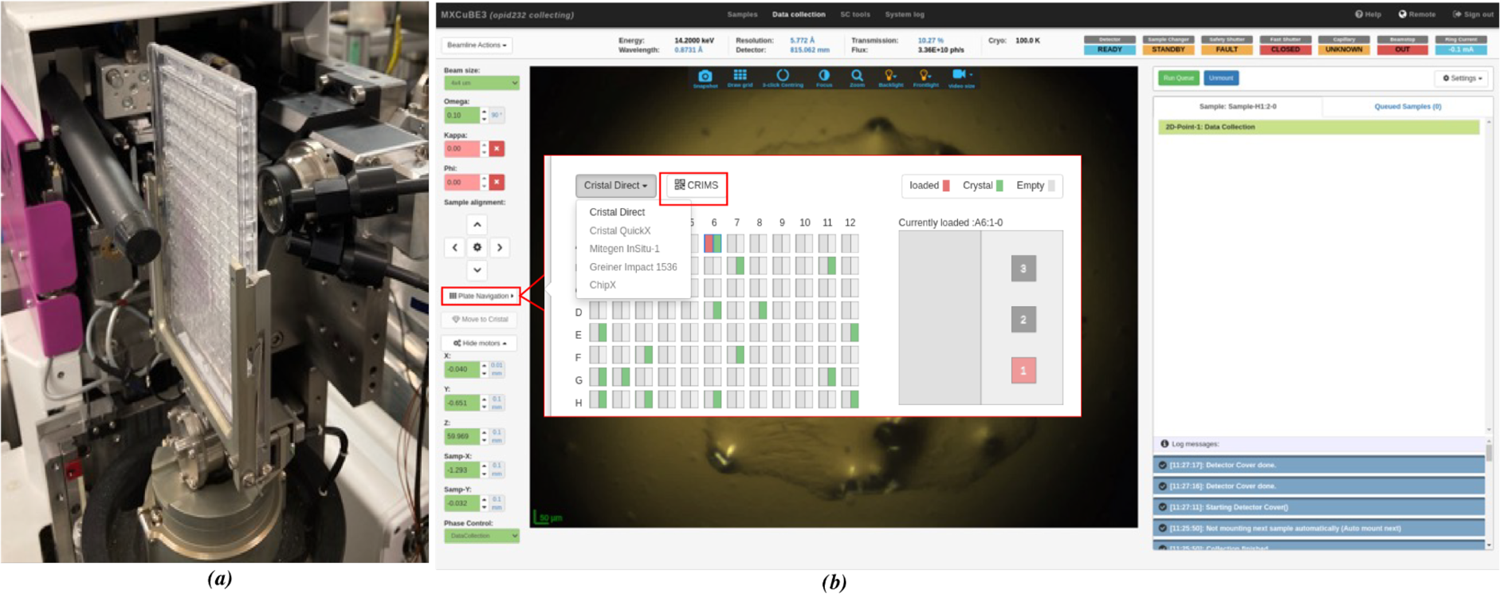
*In-situ* data collection setup with CrystalDirect plate shown as mounted on ID23-2. (*a*) CrystalDirect plate placed into the plate-manipulator (PM) on MD3-Up. (*b*) *MXCuBE-Web* interface to control *in-situ* data collection from plates. A plate-navigator button is highlighted in red-colour to emphasise the ease of going through different drops and the possibility to synchronise with the *CRIMS* database for importation of visual inspection scores (red inset). Additionally, a drop-down menu enlists different SB 96-well plates supported by the PM goniometer-head.

### 3.2 Experimental workflow control softwares

#### 3.2.1 MXCuBE-Web

*MXCuBE-Web* is the latest generation of the data acquisition software *MXCuBE* (Macromolecular Xtallography Customized Beamline Environment) (Oscarsson *et al*., 2019) – a cross-facilities collaborative project in Europe. To facilitate *in-situ* data collection, *MXCuBE-Web* has been enriched with new functionalities. Our implementation (described in Supplementary Text) considers the PM as a sample changer itself; each well is designated as a container and each drop represents a sample. The visual scores on the drops inside a 96-well plate, executed by users in their home labs, can easily be imported via a .XML file to *MXCuBE-Web*. Thus, users can directly navigate to the drops of interest with the corresponding crystals and subsequently collect diffraction data (**Figure 2**) on a single crystal or multiple ones using different collection strategies available through *MXCuBE-Web*. In this work, we have simplified visual score import by exploiting the unique plate number attributed by the High-throughput Crystallisation (HTX) platform at EMBL-Grenoble. This allows direct communication with the Crystallisation Information Management System (*CRIMS*) to retrieve the drop pointed out as containing crystals of interest. *CRIMS* is a web-based laboratory information system that provides automated communication between crystallisation and synchrotron data collection facilities, enabling uninterrupted information flow over the whole sample life-cycle from pure protein to diffraction data (Healey *et al*., 2021).

#### 3.3.3 Micro-diffractometer software

The MD3-Up control software runs on a Windows PC and is based on the *JLib* library (EMBLEM Technology Transfer GmbH, Heidelberg, Germany; http://software.emblem.de). The MD3-Up can be controlled using a graphical user interface (GUI) or remotely by a socket server using the Exporter protocol of *JLib*. The on-axis prosilica video camera of the MD3-Up is read using the *LIMA* generic library for high-throughput image acquisition (Homs *et al*., 2011). Within the MD3-Up control software a dedicated menu allows the user to navigate on the entire plate. Each type of plate benefits from its own layout for simple navigation.

## 4. Materials and methods

### 4.1. Micro-crystallization of rat autotaxin

Crystallisation was performed using the HTX platform at the EMBL Grenoble, in CrystalDirect (CD) plates. Crystals were obtained using an adapted protocol of the conditions previously described for r-ATX-beta (Eymery *et al*., 2023). The r-ATX-beta protein solution was concentrated to 3-4 mg/mL in Tris 50 mM, NaCl 150 mM at pH 8. The best diffracting microcrystals were obtained in 20-32% PEG3350, 0.1-0.4 M NH_4_I, 0.1-0.4 M NaSCN using the hanging drop method. Typically, higher concentrations of PEG3350 resulted in smaller crystals with higher density per drop. In each CD plate, 96 wells were filled with precipitant and three drops per condition were set up at different protein-precipitant ratios (1:1, 1:2, and 1:3). Promising drops (**Figure S1**) with high microcrystal density were ranked for data collection after automated visible and UV light imaging, and visual inspection scores were registered in the *CRIMS* database.

### 4.2. *In-situ* serial crystallography (iSX) data collection from CrystalDirect plate

At ID23-2, data collection is conducted via *MXCuBE-Web* application (Oscarsson *et al*., 2019) and results/outputs of data collection and automated processing are displayed in the data-management system, *EXI-ISPyB* (Delageniere *et al*., 2011). *MXCuBE-Web* has been adapted to support *in-situ* (RT) data collection from plates. *MXCuBE-Web* in “plate mode” enables users to import visual inspection scores for each CrystalDirect plate, previously conducted at the EMBL-Grenoble HTX facility by synchronising with the *CRIMS* database (Cornaciu *et al*., 2021; Healey *et al*., 2021) (**Figure 2b**). We collected ∼5.7 million diffraction frames from 61 drops, across 2 CrystalDirect plates using MeshScan (Zander *et al*., 2015), a modular automated ESRF workflow system (EWOKS) routine available through *MXCuBE-Web*. Each drop was raster scanned with a 4 ✕ 4 µm^2^ X-ray beam at 14.2 keV photon energy, a flux of 7×10^11^ ph/sec (i.e., 10% of full flux) and 6 ms exposure per frame without any rotation. This *in-situ* serial data collection strategy yielded ∼100,000 (average) still diffraction frames per drop in ∼10 min, ensuring the diffraction weighted dose to be ∼35 kGy/diffraction frame, as calculated using *RADDOSE-3D* (Zeldin *et al*., 2013).

### 4.3. Cryogenic serial (cryo-SSX) data collection on Autotaxin

Diffraction data were collected at ID23-2 at the European Synchrotron Radiation Facility (ESRF), Grenoble, France (Nanao *et al*., 2022). Measurements from r-ATX microcrystals were carried out under cryogenic conditions (100 K) with 4 × 4 μm^2^ microfocus beam at 14.2 keV (wavelength at 0.8731 Å). Entire crystallisation drops were harvested and raster scanned with X-ray exposure times of 6 ms/image to identify positions corresponding to well diffracting crystals. In a second step, 10° wedges consisting of 100 frames with 0.1° oscillation were collected with 20 ms/frame exposure time from each of the identified positions at a speed of 5°/sec using the *MeshAndCollect* automated multi-crystal data collection workflow (EWOKS) (Zander *et al*., 2015). This strategy ensured the diffraction weighted dose to be 3 MGy/sweep as calculated by *RADDOSE-3D* (Zeldin *et al*., 2013). If dose allowed, 2-3 sweeps of 10° wedges were collected from each identified crystal. Each 10° wedge takes typically 2 sec.

## 5. Data-processing, structure determination and refinement

### iSX data processing

The still diffraction frames, collected at room temperature, were processed using *crystFEL* v.0.10.1 (White *et al*., 2012) distributed within *SBGrid* (Morin *et al*., 2013). Peak-finding was done using the *peakfinder8* algorithm (Barty *et al*., 2014). The indexing step was performed with *indexamajig* combined with *xgandalf* (Gevorkov *et al*., 2019) and *mosflm* (Battye *et al*., 2011). Data were merged at the resolution cutoff of 3.0 Å using *partialator* with *unity* model (*CrystFEL*). The statistics reported (**Table 1**) have been calculated using *check_hkl* and *compare_hkl* (*CrystFEL*)*. TRUNCATE* (French & Wilson, 1978) from *ccp4i* (Agirre *et al*., 2023) has been used to convert Intensities (I) to Structure Factors (SF). The structure was determined by molecular replacement using *PHASER* (Read, 2001) from *phenix* (Adams *et al*., 2010) with the PDB model 4ZGA (Stein *et al*., 2015) as a reference model homologous at 92.2% in sequence calculated with *SIM* (Duret *et al*., 1996) from the expasy.org web server. The reference model was prepared using *PDBSET* (*ccp4i*). Refinement was carried out using *phenix.refine* (Afonine *et al*., 2012) for reciprocal space and *Coot* (Emsley & Cowtan, 2004) for real space, yielding R_free_/R_work_ values of 0.28/0.22 respectively.

**Table 1.** Data-collection and refinement statistics for rat autotaxin (r-ATX). Values in parentheses represent the highest resolution shell.

|  | *r-ATX - iSX* | *r-ATX - Cryo-SSX* |
| --- | --- | --- |
| Data availability | [doi:10.15151/ESRF-DC-1565254585](https://doi.org/10.15151/ESRF-DC-1565254585) | [doi:10.15151/ESRF-DC-1562957625](https://doi.org/10.15151/ESRF-DC-1562957625) |
| PDB-ID | 9EU5 | 9ENT |
| **Data statistics** |  |  |
| Space group | *P*1 | *P*1 |
| *a*, *b*, *c* (Å) | 54.51, 61.19, 65.001 | 53.51, 61.19, 65.01 |
| $\alpha$, $\beta$, $\gamma$ (°) | 102.568, 99.128, 93.67 | 102.568, 99.128, 93.67 |
| # Diffraction frames collected | 5, 557, 065 | 934 x 100 |
| # Indexed frames merged | 29,553 | 91 x 100 |
| Resolution range (Å) | 62.448 - 3.0 (3.14 - 3.04) | 37.51 - 2.2 (2.27 - 2.2) |
| Total No. of reflections | 2,139,506 (116,235) | 270,477 (19,435) |
| # unique reflections | 32,245 (2,669) | 34,797 (2,597) |
| Completeness (%) | 99.96 (99.89) | 99.96 (99.89) |
| Multiplicity | 66.35 (43.6) | 7.77 (7.48) |
| $\langle I/\sigma(I) \rangle$ | 4.31 (1.79) | 4.45 (0.97) |
| CC<sub>1/2</sub> ‡ | 0.87 (0.56) | 0.952 (0.40) |
| **Refinement statistics** |  |  |
| Resolution range (Å) | 62.45-3.00 | 37.51-2.2 |
| # unique reflections/test set | 16,129/825 | 39,724/1986 |
| $R_{free}/R_{work}$ ◆ | 0.2809/0.2245 | 0.2357/0.1938 |
| No. atoms | 6688 | 6968 |
| Protein | 6354 | 6424 |
| Others | 131 | 95 |
| Water | 203 | 449 |
| **B-factors** |  |  |
| Wilson plot | 59.34 | 38.32 |
| Total | 55.31 | 47.33 |
| Protein | 55.17 | 46.93 |
| Water | 44.91 | 49.7 |
| Others | 78.11 | 63.18 |
| **Root-mean-square deviation ◇** |  |  |
| Bond lengths (Å) | 0.003 | 0.003 |
| Bond angles (°) | 0.588 | 0.643 |
| Ramachandran |  |  |
| Favoured (%) | 95.52 | 96.02 |
| Outliers (number) | 0 | 0 |

‡ CC<sub>1/2</sub> values that are significant at the 0.1% level are marked by an asterisk. ◆ Crystallographic R factor, $R_{work} = \sum ||F_o| - |F_c|| / \sum |F_o|$ where $F_o$ and $F_c$ are the observed and calculated structure factors, respectively. $R_{free}$ is $R_{work}$ calculated for a randomly selected subset of 5% of the data for the r-ATX crystals, not used in the refinement procedure. ◇ Root-mean-square deviations from the standard values for bond lengths and angles.

### Cryo-SSX data processing

Initial diffraction patterns, collected under cryogenic condition at ID23-2, were processed using XDS (Kabsch, 2010), followed by real-time automated selection and merging of statistically equivalent datasets with XSCALE using the *sxdm* tool (Basu *et al*., 2019). Datasets were selected using a combination of multiple criteria, including ISa cut-off of 3.0, unit-cell, and pair-CC based hierarchical clustering. This allowed us to quickly identify 91 statistically equivalent mini-datasets (i.e., 10° wedges), which were then scaled and merged using *XSCALE*. Thus, a reflection file (MTZ format) was produced for further structural analysis. All the structures were solved by molecular replacement (MR) using *PHASER* (McCoy, 2006) with the PDB model 4ZGA (Stein *et al*., 2015) as a search model. The resolution cut-off was determined to be 2.2 Å using a combined metric of CC_1/2_ of ≥ 0.30 and I/σ(I) of ≥ 1.0 (Karplus & Diederichs, 2012). Structure refinement was carried out iteratively with *phenix.refine* (Afonine *et al*., 2012) and *Coot* (Emsley & Cowtan, 2004). *MolProbity* (Williams *et al*., 2018) was used to assess the quality of the structure and the data collection and refinement statistics are summarised in **Table 1**.

### 5.1 Space group and unit cell determination

As r-ATX was crystallised in a triclinic lattice, unit-cell determination on still diffraction frames obtained from millions of different microcrystals was challenging. Primarily, indexing on each drop containing a reasonably high number of crystal hits was executed with the *indexamajig* program within *CrystFEL* using the *mosflm* and *xgandalf* algorithms. The diffraction frames, indexed in triclinic lattice, were clustered in multiple unit-cell parameters. Using these initial results, we were able to find three “crystals populations’’, out of which, two populations appeared more isomorphous than the 3rd one.

Thus, we determined two sets of cell parameters (cell-I) *a, b, c* (Å): 54.16, 62.66, 65.55 and *α, β, γ* (°): 77.65, 81.24, 93.80 and (cell-II) *a, b, c* (Å): 53.5, 61.19, 65.01 and *α, β, γ* (°): 102.56, 99.12, 93.67. We used these cell parameters to repeat the indexing step. The indexing, performed with cell-II parameters, had a better success rate when compared to indexing executed with cell-I. We therefore focused on the results of indexing obtained with cell-II parameters. Subsequently, we performed indexing with an increased unit-cell tolerance parameters, 12% on *a*, b*, c\** axes and 5% on angles. This strategy gave us the best results with 29,553 indexed frames from a total of 5,557,065 diffraction frames collected. The final indexing results appeared to be slightly non-isomorphic, which was investigated using the *k-means* analysis (**Figure 3**) to determine the degree of heterogeneity. **Figure 3** quantified that the suspected two populations of crystal cells were actually very close to each other with an Euclidean vector distance of 3.28 Å, which corresponds to a negligible difference of 0.09% in unit-cell volume. To complete the data analysis, we further visualised Bragg reflections in 3D using *phenix data 3D viewer* tool and observed no anisotropicity due to preferred orientation, which would normally be expected from r-ATX crystal morphology (**Figure S2)**.

**Figure 3.**
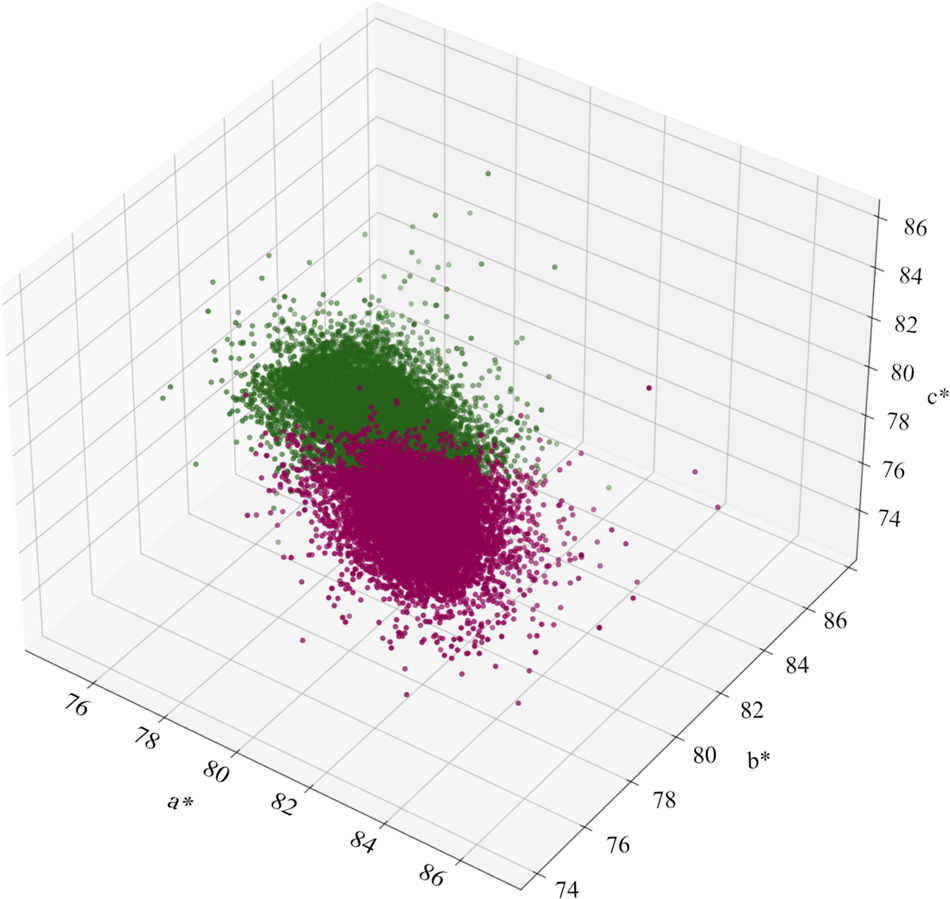
Final distribution of crystal unit cell dimensions from the *in-situ* dataset. Each dot represents a crystal lattice. For visualisation, unit-cells were orthogonalised in reciprocal space. In 3D cartesian coordinate, each dot is associated with *a*, b*, c\** vector cell along x, y, z axes respectively, with units in 1/Å. *a*, b*, c\** have been calculated using the cosine law formula 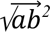, 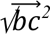, 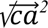. In green and purple are presented each population separated using the *k-means* method.

### 5.2 Merging and resolution cut-off

Indexed frames were merged using *partialator* in *CrystFEL*. The resolution of the merged dataset was truncated at 3.0 Å considering the behaviour of R_split_, CC*, CC and *<I/σ* (*I*)*>*. All the data processed for the *á posteriori* analysis has been truncated at 3.0 Å resolution to be directly comparable with our reference data set. Our *conservative* resolution cut-off was decided based on the heterogeneity due to different crystals and variable sizes and diffraction quality. The variability within the sample has an impact on the proper estimation of the error modelling of the intensity measurements. We took this information into account and decided to cut the data at a pretty high *<I/σ*(*I*)*>* value (Wlodawer *et al*., 2008; Evans & Murshudov, 2013).

### 5.3 Optimise future experiment by *a posteriori* analysis

An *a-posteriori* analysis of our data was executed to determine an optimal beamtime session for a large triclinic structure. To this end, we ranked the drops in descending order based on the number of indexed frames, followed by merging them. Thus, we estimated the minimum number of drops required to obtain a complete dataset for r-ATX in the triclinic lattice. We prepared 10 datasets by merging indexed frames from the “best” 5, 6, 8, 10, 15, 17, 18, 20, 25 and 30 drops, respectively. Using this method, we determined that after iSX data collection on 10 drops, we had a usable dataset and after 17 drops, the CC* began to converge close to the value obtained from all-data, comprising 61 drops (**Figure 4**). Molecular replacement (MR) was performed for all sub-datasets using the same model as for the original phasing. Subsequently, the MR solutions were compared to visually inspect the electron density interpretability from each of the datasets (**Figure 5**). From the best_10 dataset onwards, manual model building in real space was clearly possible. This corresponds to the effective data collection time of 1.5 hours across 10 drops.

**Figure 4.**
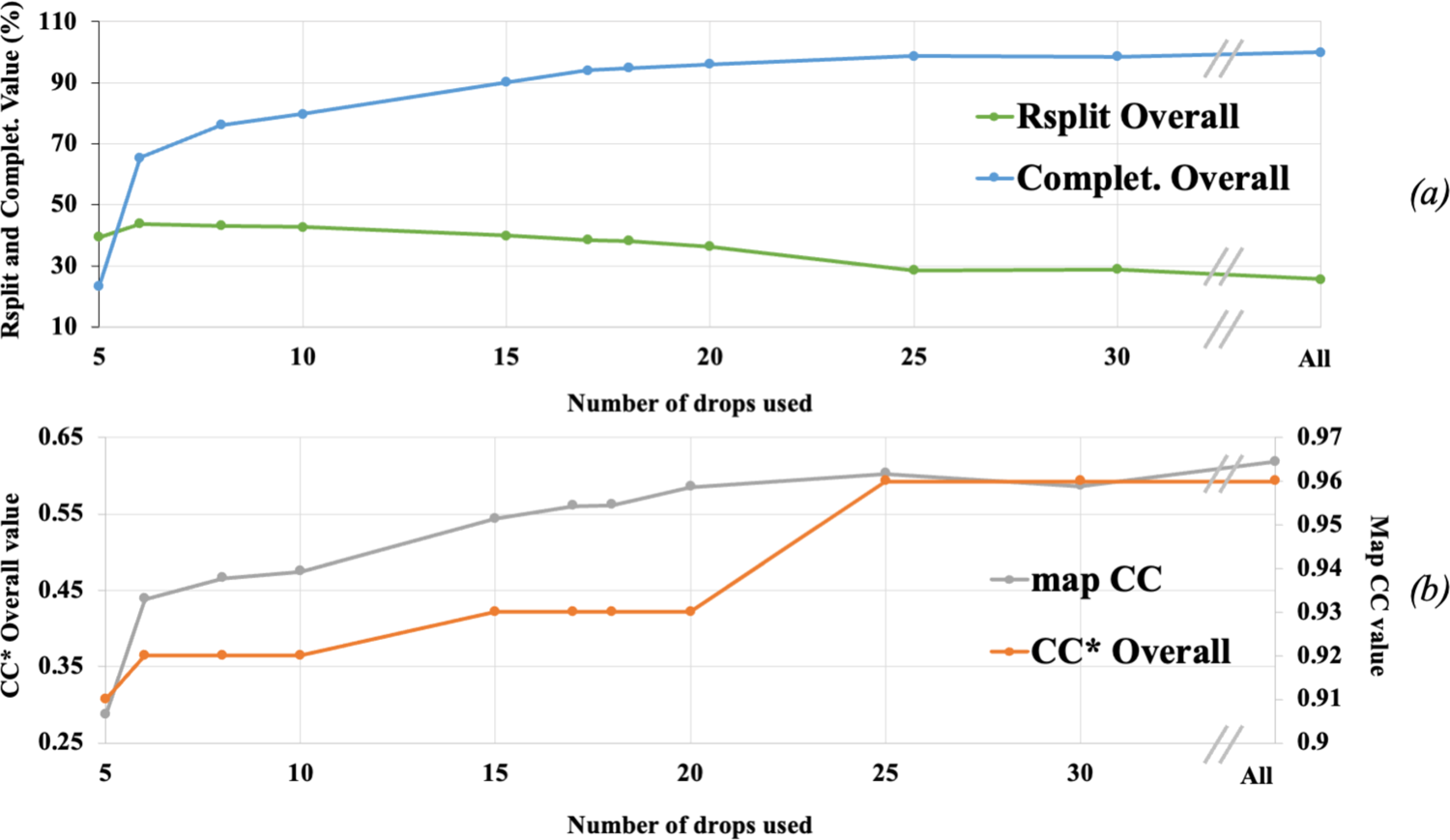
Merging statistics of iSX datasets plotted against the number of drops for each dataset, from 5 to 30 drops and comparison of each dataset used for refinement. Panel *(a)*: In green: R_split_ Overall (%) and in cyan: Completeness (%). Panel *(b)*: In orange CC* Overall (right axes) and in grey map correlation (left axes) between the first 2mFo-DFc electron density map obtained after the molecular replacement and the final refined “All”, implying 61 drops, 2mFo-DFc electron density map.

**Figure 5.**
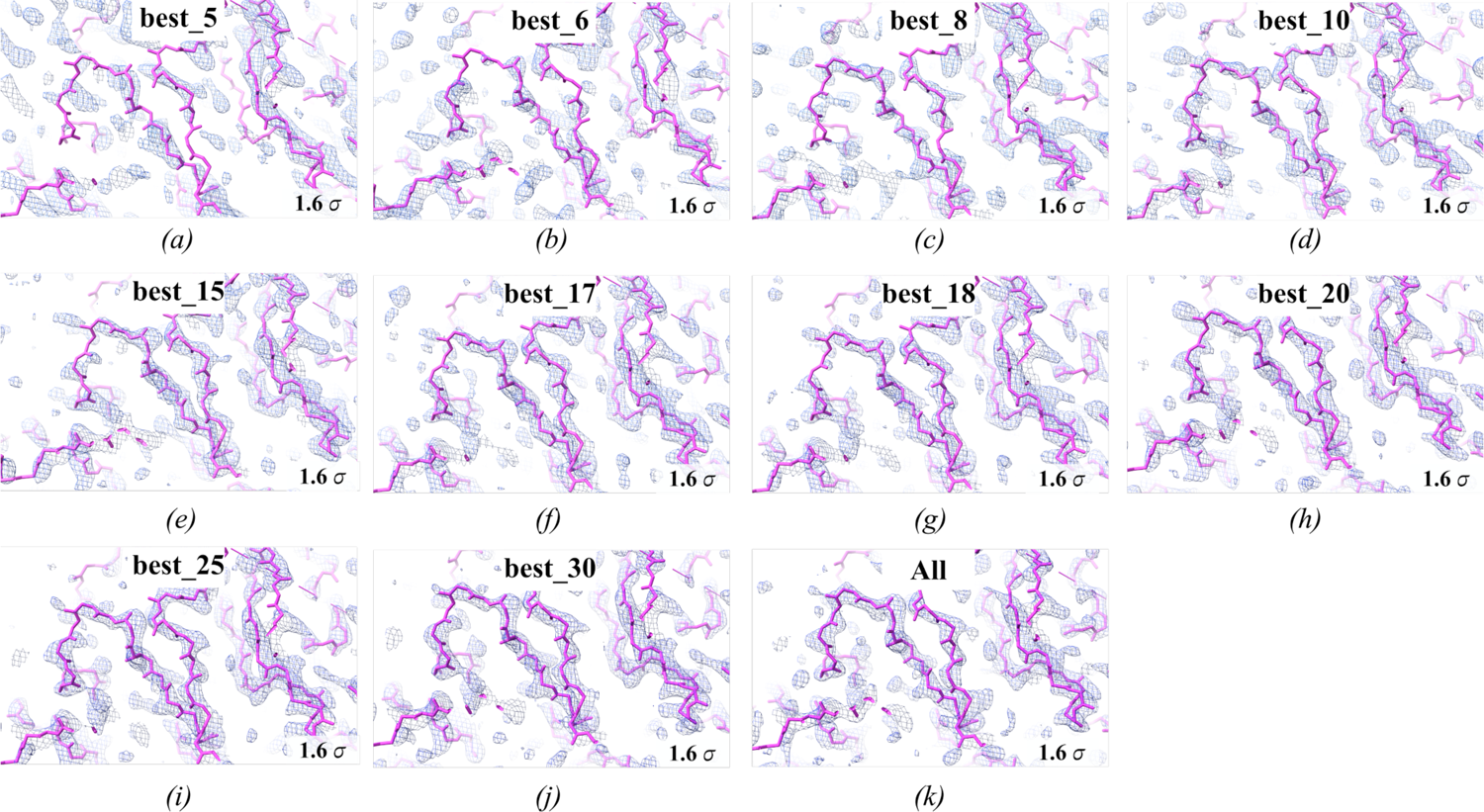
*In-situ* serial crystallography (iSX) structure of r-ATX solved in *P*1 space group. Comparison of first 2Fo–Fc electron density map at 1.6 σ level from molecular replacement result produced by *phaser* and displayed as a blue mesh overlaid onto r-ATX C⍺ model (pink sticks). Each panel represents respectively *(a)*: best_5, *(b)*: best_6 *(c)*: best_8, *(d)*: best_10, *(e)*: best_15, *(f)*: best_17, *(g)*: best_18, *(h)*: best_20, *(i)*: best_25, *(j)*: best_30 and *(k)*: all-data: 61. Qualitatively the electron density is easily interpretable from the *(d)*: best_10 onwards.

### 5.4 Structure comparison between iSX and cryo-SSX

We compared the iSX and cryo-SSX structures of r-ATX by superimposition and RMSD calculation using *pymol* (DeLano, 2002) (**Figure 6 and S3**). Upon structural comparison between iSX and cryo-SSX models of r-ATX, we did not observe any differences at the active site. However, both – iSX and cryo-SSX structures reveal an oxysterol moiety bound in the allosteric tunnel (**Figure 6**). We compared the relative B-factor-change (∂B_relative_) between the iSX and cryo-SSX structures of r-ATX. For each model, B-factors were averaged over all residues followed by subtraction of B-factor of cryo-SSX model from *in-situ* model for each residue. We applied a fixed positive offset over all the values to have the lowest value at zero. We then used *pymol* to represent the B-factor difference combining putty and rainbow colour representation (**Figure S3**) (DeLano, 2002). We calculated the overall RMSD between the two models resulting in a value of 1.34 Å **(Figure S3)**. We observed the largest RMSD value for small domains (**Figure S3a**) outside of the protein core in regions that are located along the solvent channel considering the crystal packing (**Figure S3b)**. The larger volume in such a location allows a wider distribution of possible loop conformations. The B-factor difference between *in-situ* and cryo structures follows the same trend as the RMSD. The higher B-factor differences are colocalising in more flexible domains at the protein surface in the crystal contact regions.

**Figure 6.**
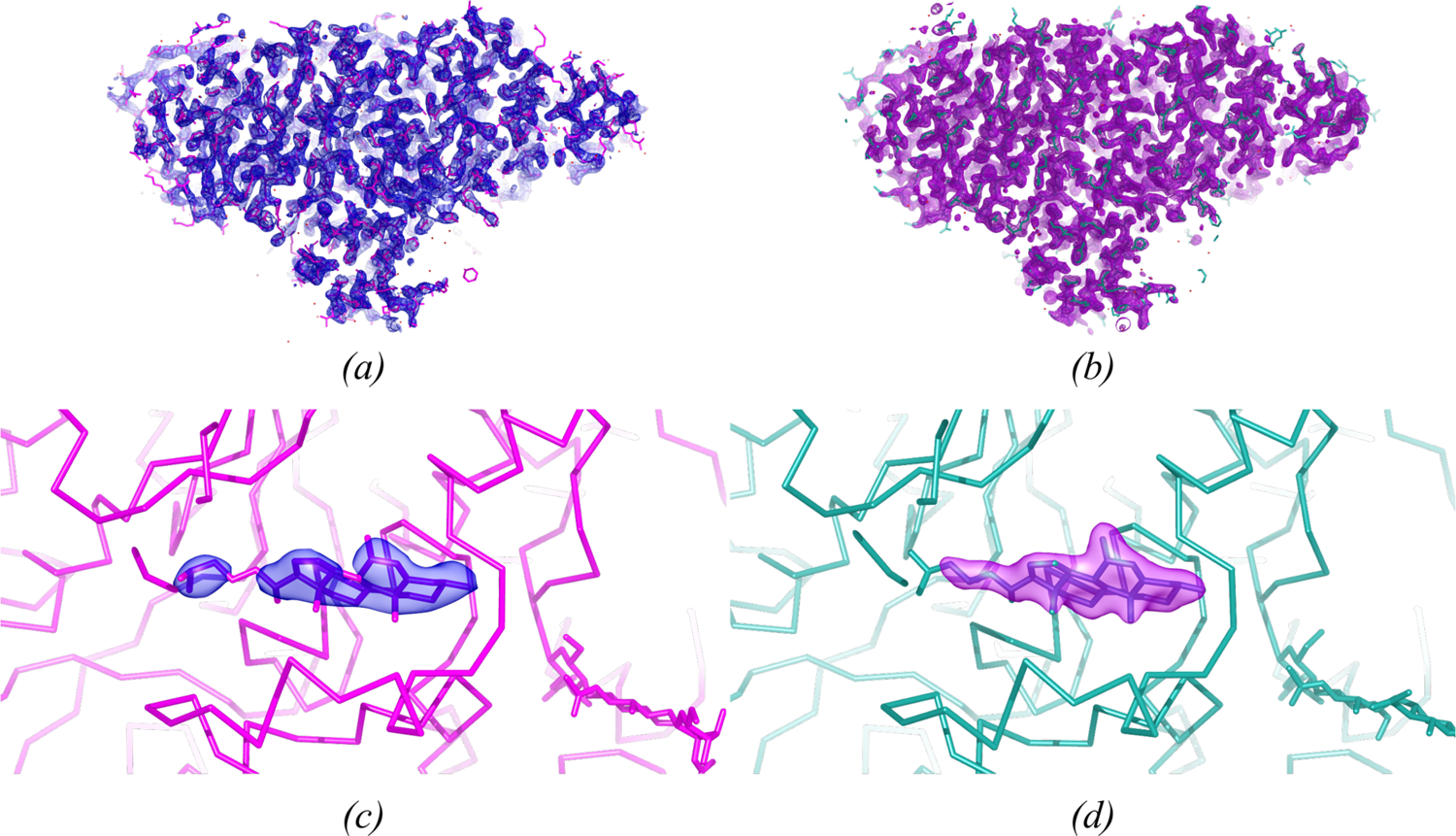
Serial crystallography structure of r-ATX in *P*1 space group. *(a)* and *(c)* displays the *in-situ* determined structure, *(b)* and *(d)* displays structure determined from serial cryo data collection. *(a)* overall 2Fo–Fc electron density map at 1.2 σ level in blue overlaid onto r-ATX model in magenta sticks representation. *(c)* highlights the 2Fo–Fc electron density map coloured in blue for the oxysterol moiety bound in the allosteric tunnel of the r-ATX model shown in magenta. *(b)* overall 2Fo–Fc electron density map at 1.2 σ level in purple overlaid onto r-ATX model in ocean green sticks representation. *(d)* highlights the 2Fo–Fc electron density map coloured in purple for the oxysterol moiety bound in the allosteric tunnel of r-ATX model shown in ocean green.

## 6. Discussion

This work combines the advantages of conventional high throughput crystallisation in plates with the rapidity of serial crystallography and demonstrates the strength and efficacy of iSX, even for difficult cases. Our iSX approach relies on easy and rapid raster scanning at any modern microfocus beamline, such as ID23-2, that is equipped with a state-of-the-art micro-diffractometer and new generation X-ray detector with fast data acquisition capabilities. The iSX strategy may be envisaged to help users in cases where a novel target without any prior information of the diffraction quality is obtained, potentially allowing the acquisition of a complete dataset. An alternative approach that could have been applied is the *Mesh & Collect* protocol, which has the advantages of including small rotations, but has the disadvantages of longer data collection times and general impracticality at room temperature. However, such a strategy can only be successful at room temperature if very low doses are used for both diffractive mapping and mini-rotation collections. One could reduce the dose significantly, but depending on how the dose budget is spent, this could be extremely difficult to process with conventional MX software. Interestingly, upon visual inspection of Bragg reflections from iSX dataset in 3D reciprocal volume, data appeared to be complete up to 3 Å and preferred orientation was not observed (**Figure S2**), even though the microcrystals were plate shaped. Such crystal morphology would normally be expected to have an orientation bias. Importantly, most of the challenging large protein complexes (e.g., r-ATX) are expressed in mammalian cells, typically yielding a few mg/mL purified protein concentration. Moreover, scaling up the protein production for higher eukaryotic expression systems can be a very expensive and time-consuming process. Microcrystallization for such cases through batch methods is not suitable; that seems to be a bottleneck in expanding serial crystallography (Beale *et al*., 2019) on the many challenging medically relevant targets (e.g., membrane proteins). Thus, a 96-well crystallisation plate might be able to serve as a more efficient fixed target sample delivery tool for serial crystallography experiments on such targets. Our iSX approach is easy, rapid, and potentially reduces preferred orientation issues because crystals are not deposited on solid supports. Our work also shows that iSX can tackle even the most challenging targets, *i.e* small crystals from large proteins that do not diffract to high resolution.

We collected still images by raster scanning an X-ray microbeam without rotation. This implied data processing with serial crystallography software, e.g., *CrystFEL*. We deliberately chose a set of parameters for hit finding that were tolerant, resulting in a large number of images considered as hits (more false positive), many of which were not possible to index in subsequent steps. As we did not benefit from any rotation information, the indexing has been quite challenging for the triclinic lattice. However, we managed to iteratively refine our cell parameters from an initial heterogeneous distribution of cell parameters. Subsequently, 29,550 indexed diffraction frames were merged using *partialator* within *CrystFEL*. We determined the *in-situ* r-ATX structure in the *P*1 space group using molecular replacement, followed by refinement in PHENIX.

In addition, we determined the r-ATX structure using cryo-SSX, by collecting 10° wedges on many micro-crystals via our *Mesh & Collect* strategy. This served as a reference structure that was used to compare with the iSX model. Superimposition of the two models showed no large overall structural differences (**Figure S3b**), except variations that occurred in crystal-contact regions. This observation supports other studies, revealing that room-temperature structures are more flexible and dynamic, potentially representing a more physiological state (Fraser *et al*., 2011). Furthermore, we compared the B-factor variation per residue between the two models and showed that the variations are again in the region where the crystal packing is less constrained (**Figure S3c**). As a result, this demonstrates that structures determined via iSX are at least as informative as those resulting from cryogenic temperature and may provide structural insights into flexible domains or regions.

To demonstrate the efficiency of our method, an *a-posteriori* data analysis was performed aimed at determining the optimal data collection strategy. Thus, we deduced that we had an acceptable data set after merging indexed frames from 10 drops by evaluating the overall completeness (**Figure 4** and **Table 1**). **Figure 5** further supports the results visually through the electron density maps. The statistics and the electron densities improved by adding more indexed images and we decided to use “drops” as data collection units instead of image numbers to discuss the efficiency. This is consistent with the fact that in our experimental setup, the fast data collection is performed drop by drop. We expect users to start with the most promising drops in terms of crystal density/quality. This justifies our methodology to accumulate diffraction frames starting from the most promising drops, which contained good crystal density. Here, we focus on the effective data collection time required to obtain a minimal dataset for the structural determination of a large biomolecule of ∼100 kDa in triclinic symmetry. Thus, it can be deduced that such a large triclinic structure could be determined from a minimum of 10 crystallisation drops in less than 1.5 hrs (∼6% of a single beamtime shift) of effective data collection. In the case of iSX data collection of r-ATX at ID23-2, it is noteworthy that the sampling of the drop using raster scans is driven by the beam size, implying a very large number of diffraction frames with a microfocus beam. This had a side effect on the number of “empty” images which may yield an “artificially” low indexing rate for such a large dataset (**Table 1**).

To the best of our knowledge, this work was the first successful attempt to determine the molecular structure of a large protein in the lowest possible symmetry using an iSX approach. The ease and rapidity of our iSX strategy, we believe, will make *in-situ* data collection at ambient conditions high-throughput and universally applicable to other MX beamlines. Importantly, iSX promotes crystallisation plates as an alternative, efficient, and the most natural approach to delivering samples for serial crystallography experiments at synchrotrons.

## Acknowledgements

We are thankful for the kind gift of r-ATX expressing cell lines from the Perrakis laboratory at NKI in Amsterdam. We thank C. Mueller-Dieckmann (ESRF) for his help preparing the manuscript. We thank the high-throughput crystallisation (HTX) platform at the EMBL-Grenoble for setting up crystallisation plates. We express our gratitude to Florent Cipriani, former head of the Instrumentation team at the EMBL-Grenoble, for initiating the Plate-Manipulator project. We thank the beamline staff from the EMBL-ESRF Joint Structural Biology Group for beamtime at the European Synchrotron Radiation Facility (ESRF), Grenoble, France under proposal numbers MX-2458 and MX-2532. MC Eymery has been funded by the EMBL International PhD program. AA McCarthy and S. Basu have been funded by EMBL. N. Foos was supported by a fellowship from the career accelerator for research Infrastructure scientists (ARISE) program under Marie Skłodowska-Curie Actions grant number 945405.

## Supporting information

### S1. *MXCuBE-Web* for iSX data collection

The motorised crystal plate manipulator (PM) has been integrated into *MXCuBE-Web* as a new hardware object, which is called when the PM is the goniometer-head mounted. The hardware-object for PM is compatible with MD2S and MD3-Up diffractometers.

The PM hardware-object is based on the Sample Changer object (or Class), exploiting all the sample changer characteristics, such as: a *cell* in the sample-changer Dewar is substituted with a *row* in the 96 well-plate, a well in the plate represents a puck in the sample-changer Dewar, and a drop in the well represents the sample. It also has the mounting sample functionality, which implies a plate translational motion (horizontal and vertical) to position a drop. A plate class is organised in rows and columns – each well (Cell object) contains a drop (Drop object) and each drop could contain several crystals (Xtal object).

The PM hardware object is implemented in both QT and the latest Web versions of *MXCuBE*. In the case of *MXCuBE-2*, based on *PyQt4*, the graphical user interface (GUI) was created by combining self-contained widgets. In the *MXCuBE-Web* version, the GUI is implemented with Javascript/React-17 with the React-Bootstrap and the react-contexify libraries, displaying a more elegant context menu. In both versions, the GUI representing the 96-well plate provides a mode of interaction with the hardware object that controls the physical instrument. The code for *MXCuBE-Web* is open-source and freely available at https://github.com/mxcube/mxcubeweb. As a cross-facilities collaborative project, *MXCuBE-Web* and its plate manipulator implementation is available to other synchrotron beamlines.

---

**Figure S1.**
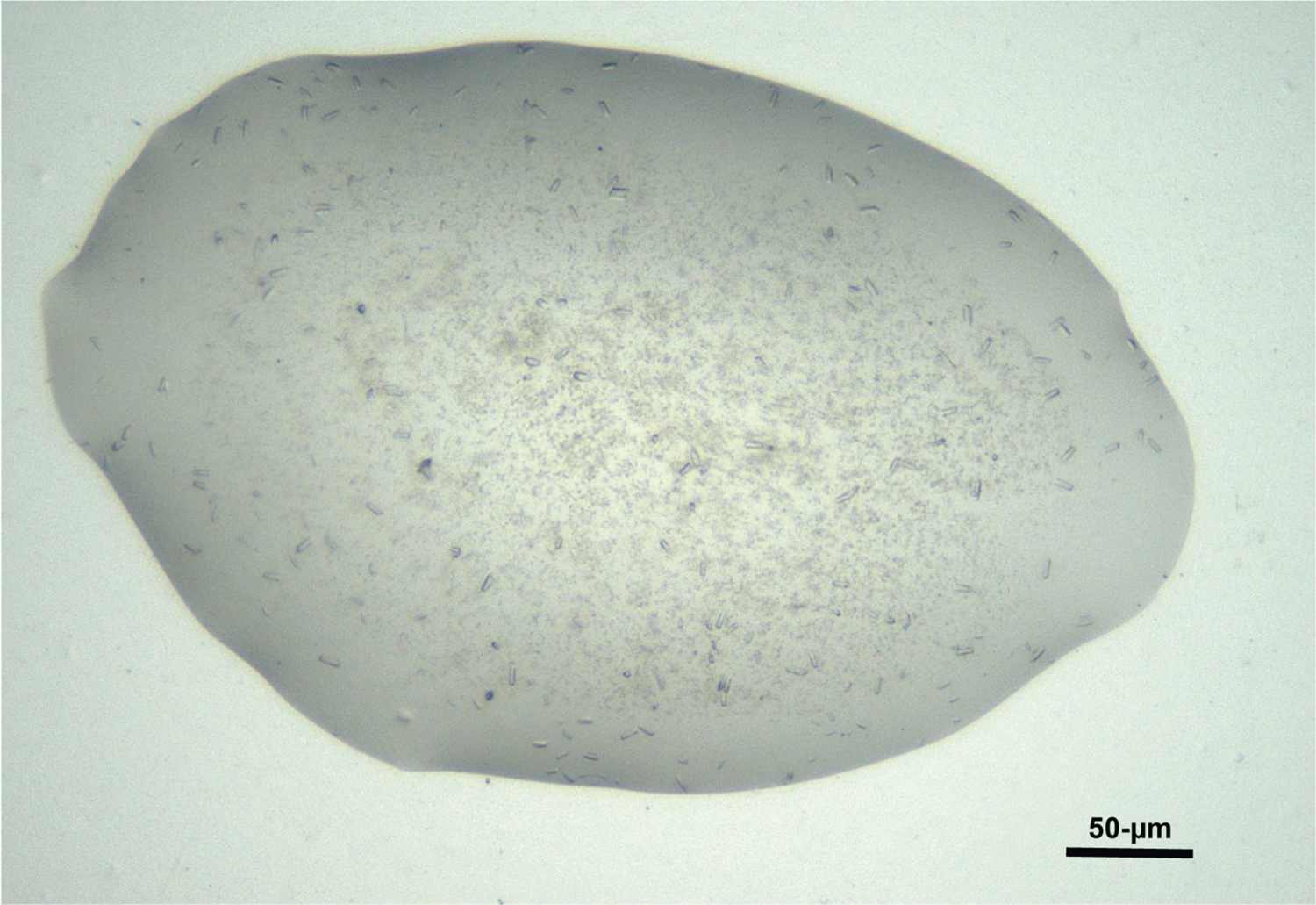
Visible microscope image of an example crystallisation drop depicting the microcrystals of r-ATX. The microcrystals are typically 5 x 5 x 10 µm^3^ in size.

**Figure S2.**
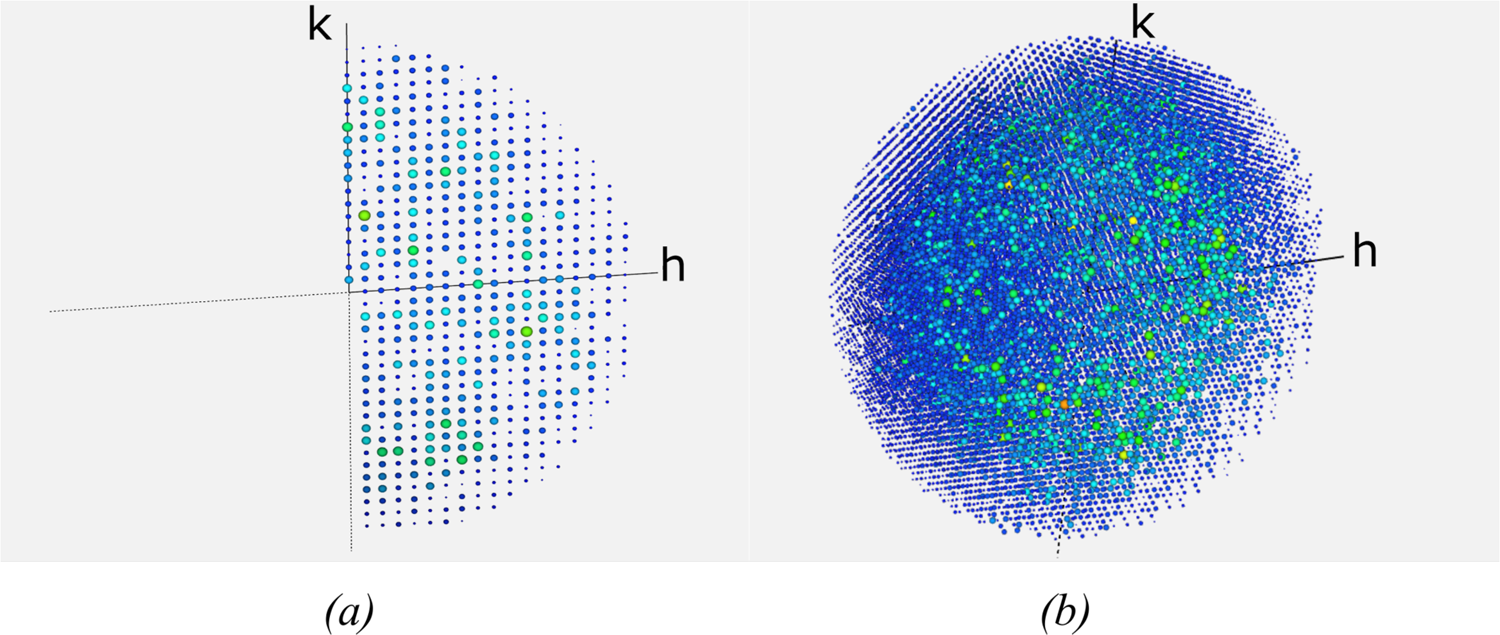
*(a)* Rainbow coloured representation of the *in-situ* dataset reflections multiplicity projection along the l axis on the h0 k0 plan. *(b)* 3D rainbow coloured representation of the *in-situ* dataset reflections multiplicity showing a complete dataset with a full sampling of the reciprocal space. The resolution cut for the representation is 3Å.

**Figure S3.**
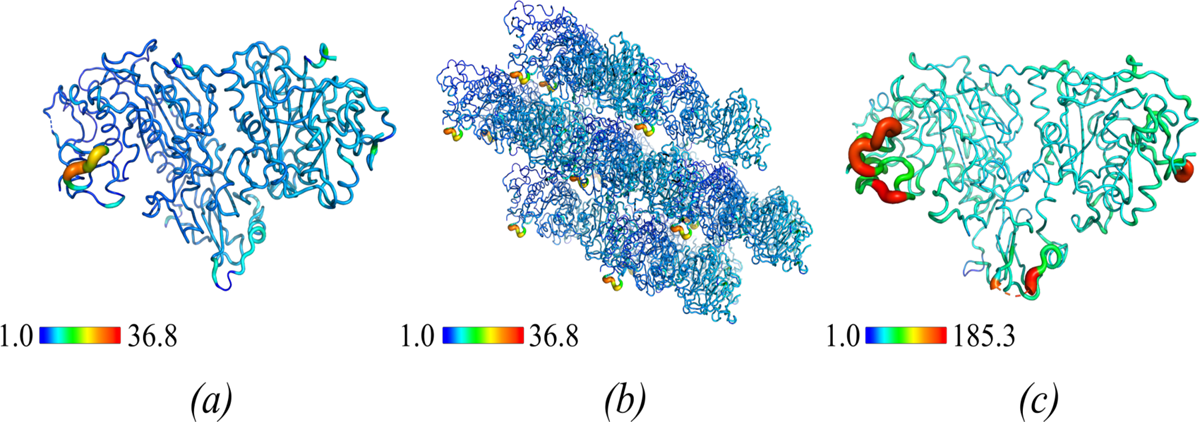
RMSD (Å) between ATX-RT and ATX-Cryo structures *(a)* and *(b)*, and comparison of B-factors (Å^2^) between ATX-RT and ATX-Cryo structures *(c)*. *(a)* Putty representation of ATX-RT model rainbow coloured. Colour and tube diameter representing the C⍺ RMSD (Å) value calculated between ATX-RT and ATX-Cryo. Loops with higher B-factor difference are shown in red and large tube diameter, and those with the lower B-factor difference are in dark blue with reduced tube diameter. *(b)* Crystal packing view. *(c)* Putty representation of the ATX-RT model rainbow coloured. Colour and tube diameter representing the C⍺ B-factors (Å^2^) differences between ATX-RT and ATX-Cryo. Loops for which the B-factor difference are higher are represented in red and large tube diameter, and those with the lower difference are in blue with reduced tube diameter.

